# Personalized expression of bitter ‘taste’ receptors in human skin

**DOI:** 10.1101/364901

**Authors:** Lauren Shaw, Corrine Mansfield, Lauren Colquitt, Cailu Lin, Jaime Ferreira, Jaime Emmetsberger, Danielle R. Reed

## Abstract

The integumentary (i.e., skin) and gustatory systems both function to protect the human body and are a first point of contact with poisons and pathogens. These systems may share a similar protective mechanism because both human taste and skin cells express mRNA for bitter ‘taste’ receptors (*TAS2Rs*). Here, we used gene-specific methods to measure mRNA from all known bitter receptor genes in adult human skin from freshly biopsied samples and from samples collected at autopsy from the Genotype-Tissue Expression project. Human skin expressed some but not all *TAS2Rs*, and for those that were expressed, the relative amounts differed markedly among individuals. For some *TAS2Rs*, mRNA abundance was related to sun exposure (*TAS2R14*, *TAS2R30*, *TAS2R42*, and *TAS2R60*), sex (*TAS2R3*, *TAS2R4*, *TAS2R8*, *TAS2R9*, *TAS2R14*, and *TAS2R60*), and age (*TAS2R5*), although these effects were not large. These findings contribute to our understanding of extraoral expression of chemosensory receptors.

## Introduction

Humans have at least five widely accepted types of taste receptors: salty, sour, sweet, bitter, and umami. The bitter receptors, called taste receptor type 2 (*T2R*), are G protein-coupled receptors that protect humans from ingesting toxins [1]. In the gustatory pathway when bitter compounds bind to a T2R protein on a taste cell, a conformational change of the protein elicits a signaling cascade. This indirectly induces the release of intracellular calcium, which leads to depolarization and neurotransmitter release, thereby activating sensory neurons that send signals to the central nervous system for bitter perception [2]. Humans have 25 bitter receptors, the T2R proteins, that are encoded by the *TAS2R* genes located on chromosomes 5, 7, and 12 (Fig 1).

**Figure 1:**
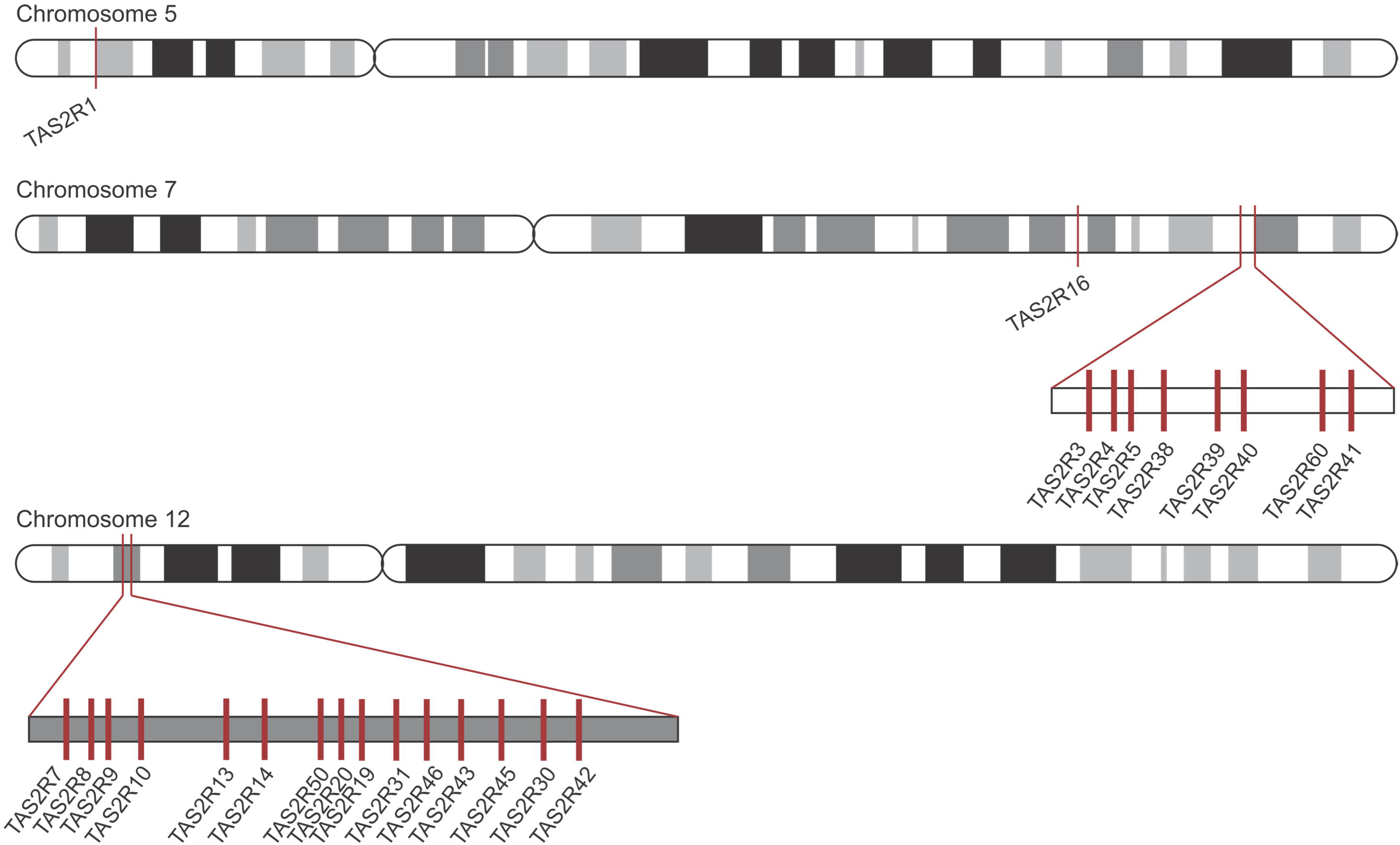
Bitter receptor locations in the human genome. The location of *TAS2R* genes on human chromosomes 5, 7, and 12 marked by red bars.

Recently, scientists have identified bitter receptors in locations of the body other than the taste cells. This expression and activation of extragustatory T2Rs will not lead to taste perception, but instead will elicit distinct cell-type-specific physiological responses. The results of several studies have demonstrated that the extraoral expression of T2Rs is involved in or regulate important biological processes germane to the nature of the tissue in which they reside. Bitter receptors have been implicated in the relaxation of smooth muscle, vasoconstriction, gut motility, bronchodilation, nutrient sensing, insulin release, and the release of the antimicrobial peptide, β-defensin [3–7]. As an example, studies performed by Lee *et al*. demonstrated that susceptibility to upper respiratory infection depends on an inborn genotype within one of these bitter receptor genes (*TAS2R38*). Gram-negative bacteria secrete a quorum-sensing molecule that is an agonist of the T2R38 receptor. People with non-functional alleles of this receptor are more susceptible to sinonasal infection because of impairments in this bactericidal pathway [8]. The broader implications of this result are that bitter receptors expressed in extraoral areas may be involved in innate immunity.

Building on this observation, we conducted a study to assess the gene expression patterns of all 25 *TAS2R* genes in skin, since it is a barrier organ and a first line of defense against invading pathogens, presenting both innate and adaptive immune functions. In addition, at least one cell type in human skin (keratinocytes) expresses olfactory receptors, which are similar to bitter taste receptors [9]. Other investigators have measured *TAS2R* mRNA expression in skin with conflicting results, perhaps owing to lack of appropriate controls against genomic DNA contamination [10, 11]. Here, we combine results from a smaller biopsy study using quantitative PCR (qPCR) and appropriate controls with a larger autopsy study using an RNA-seq method to get a more complete understanding of *TAS2R* mRNA expression patterns in human skin.

## Results

### Sample integrity

RNA and DNA were extracted from 15 whole skin samples provided by the University of Pennsylvania Department of Dermatology (Table 1) and from one fungiform taste papilla (FP) biopsy obtained from a separate donor as a representative of taste tissue. One sample (004) did not produce viable RNA (RNA integrity number equivalents = 1.0) and was eliminated from the study. Using the remaining RNA samples, cDNA was synthesized and tested for the presence of unwanted genomic DNA using the Abelson 1 (*ABL1*) gene [12]. This is a necessary step since the *TAS2R* protein-coding sequences are within single exons, and *TAS2R* primers cannot be designed to differentiate between genomic DNA and cDNA. Based on the results, three of the samples (005, 006, and 007) were unlikely to contain cDNA because they did not express this gene, and two (009 and 014) had residual genomic DNA after a second DNase treatment (S1 Fig.). These five samples were eliminated from the study.

**Table 1.**
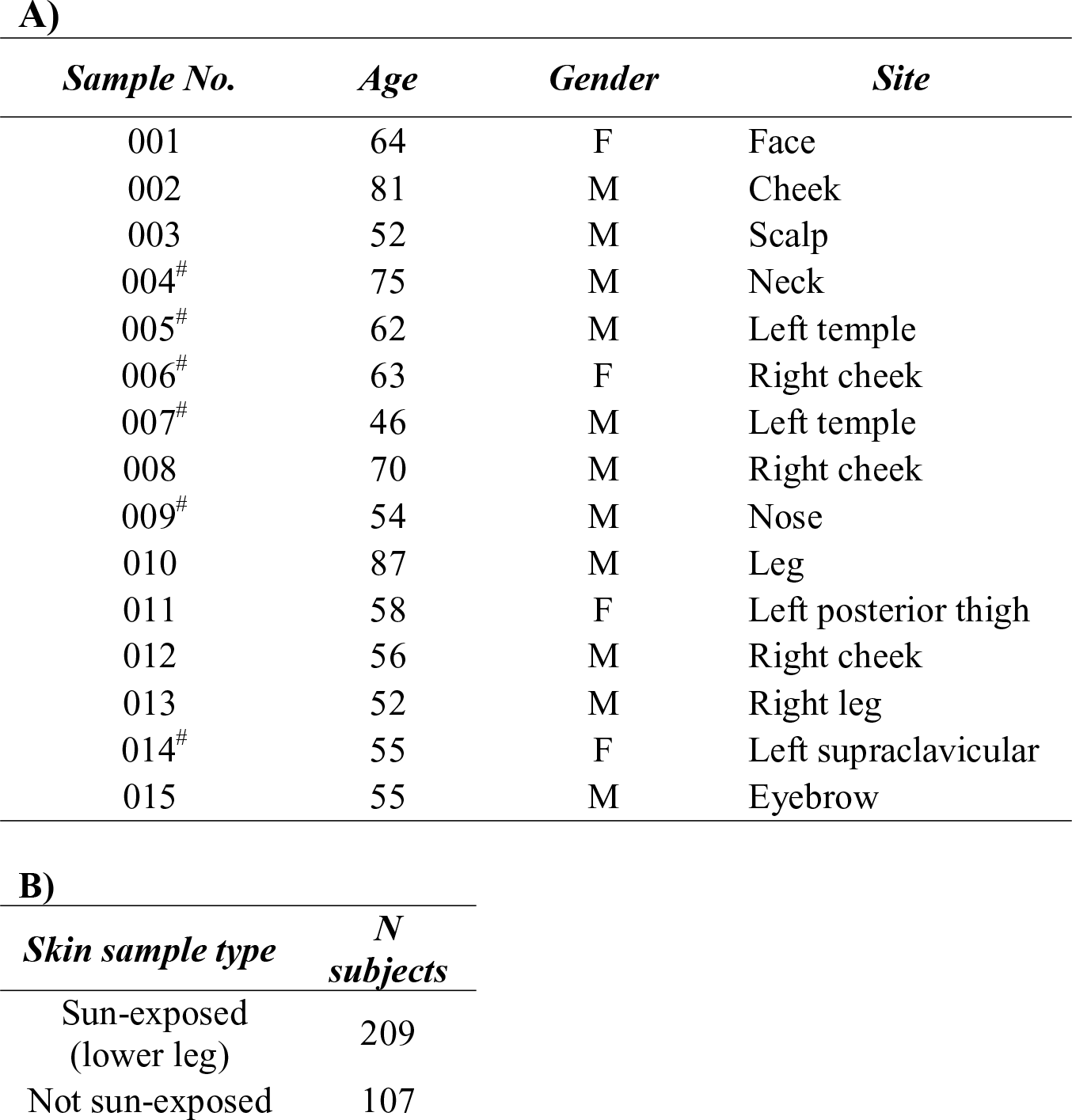
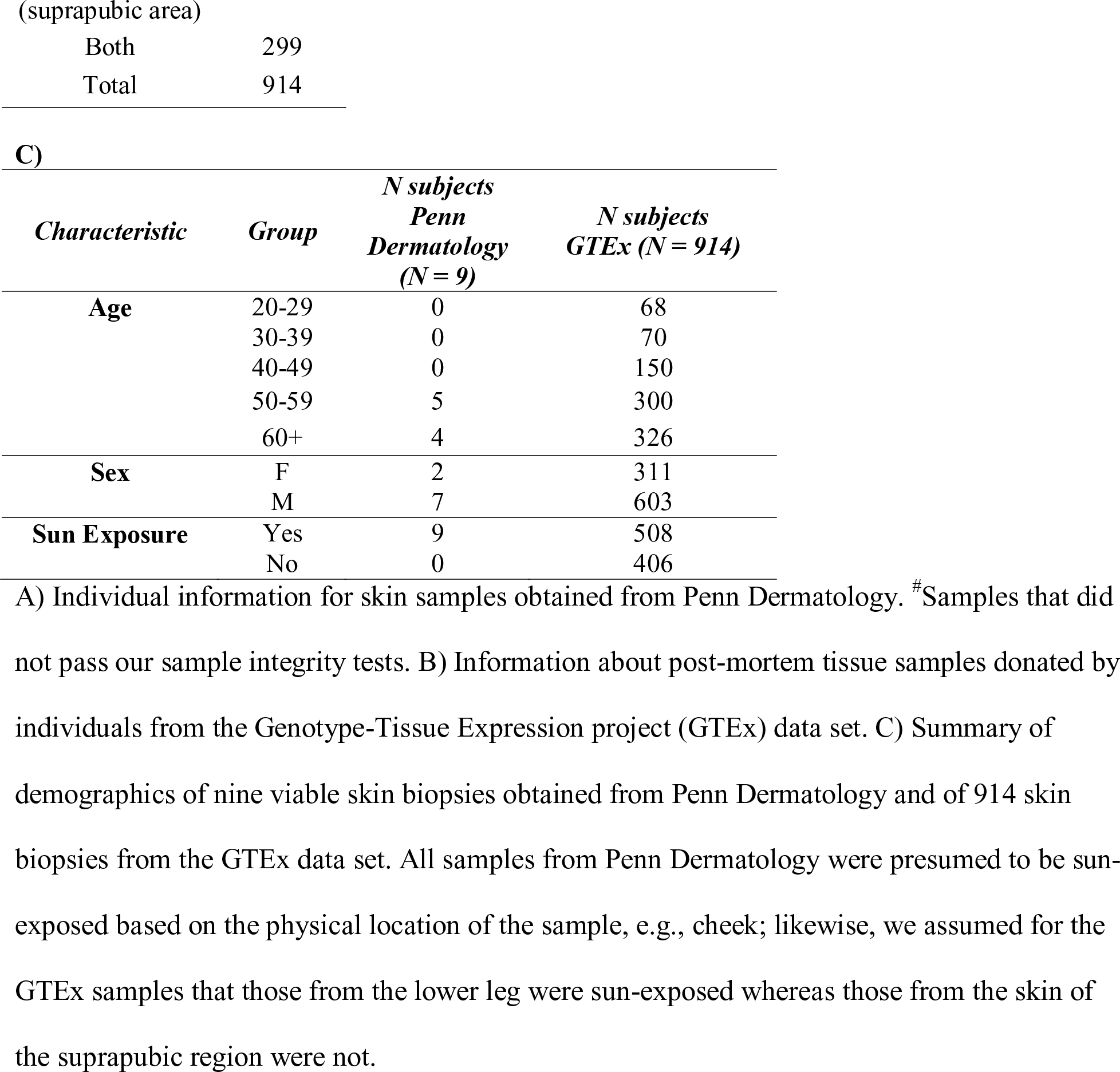
Subject characteristics.

Samples obtained from Penn Dermatology after Mohs surgery vary in size because the procedure requires surgeons to continue removing tissue until all cancerous cells are gone and only healthy tissue remains. The depth of the surgery therefore varies by individual. The samples obtained in this study consist of healthy skin that was removed to properly close the wound at the end of the procedure. Thus, each biopsy sample is unique [13]. To characterize the skin layers and cell types represented in the biopsy samples from Penn Dermatology, qPCR was performed for seven skin-layer- and cell-type-specific markers, standardized to *GAPDH*. As expected, biopsy samples differed in the relative abundance of cell-layer markers (Fig 2)[14, 15].

**Figure 2:**
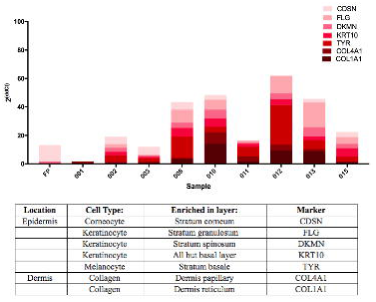
Quantification of skin-specific gene expression—qPCR results from cDNA of FP and skin samples. Data are from amplification of skin-specific markers characterized in the table [14, 15]. Data from all markers are represented in order of skin layer for each individual biopsy, with the top of the epidermis (*CDSN*) as the lightest bar section and the bottom of the dermis (*COL1A1*) as the darkest bar section. Results were standardized to the housekeeping gene *GAPDH* and expressed as 2^ΔΔCt^.

### PCR amplification

To investigate whether bitter taste receptor mRNA is expressed in human skin, PCR experiments were performed with two technical replicates for each of the 25 *TAS2R* genes (S2–S26 Figs), which were compared against two positive controls: (a) genomic DNA from skin and (b) fungiform papillae cDNA. Of the 25 *TAS2R* genes, only three showed no expression (*TAS2R1*, *7*, and *8*), 19 showed variable expression (*TAS2R3*, *4*, *5*, *9*, *13*, *14*, *16*, *20*, *31*, *38*–*43*, *45*, *46*, *50*, and *60*), and three showed universal expression (*TAS2R10*, *19*, and *30*) (Fig 3). The genomic DNA positive controls were amplified in every case; however there was variability in *TAS2R* expression in the FP, suggesting that *TAS2Rs* are expressed at low levels even in taste tissue. This low abundance may explain the variability of expression between technical replicates, as shown in S2–S26 Figs and summarized by the yellow cells in Fig 3. PCR experiments were also performed for *GNAT3*, a gene encoding for the α-subunit of the taste-associated G protein gustducin, and keratin 10 (*KRT10*), a positive epithelial marker (Fig 3 and S27–S28 Figs). *GNAT3* was detected in taste tissue, as expected, and in four skin samples (002, 003, 008, and 015), suggesting some similarity between the pathway elicited in skin and the initial steps of the gustatory pathway. As anticipated, *KRT10* was detected in FP and all skin samples. All primers are listed in Table 2.

**Figure 3:**
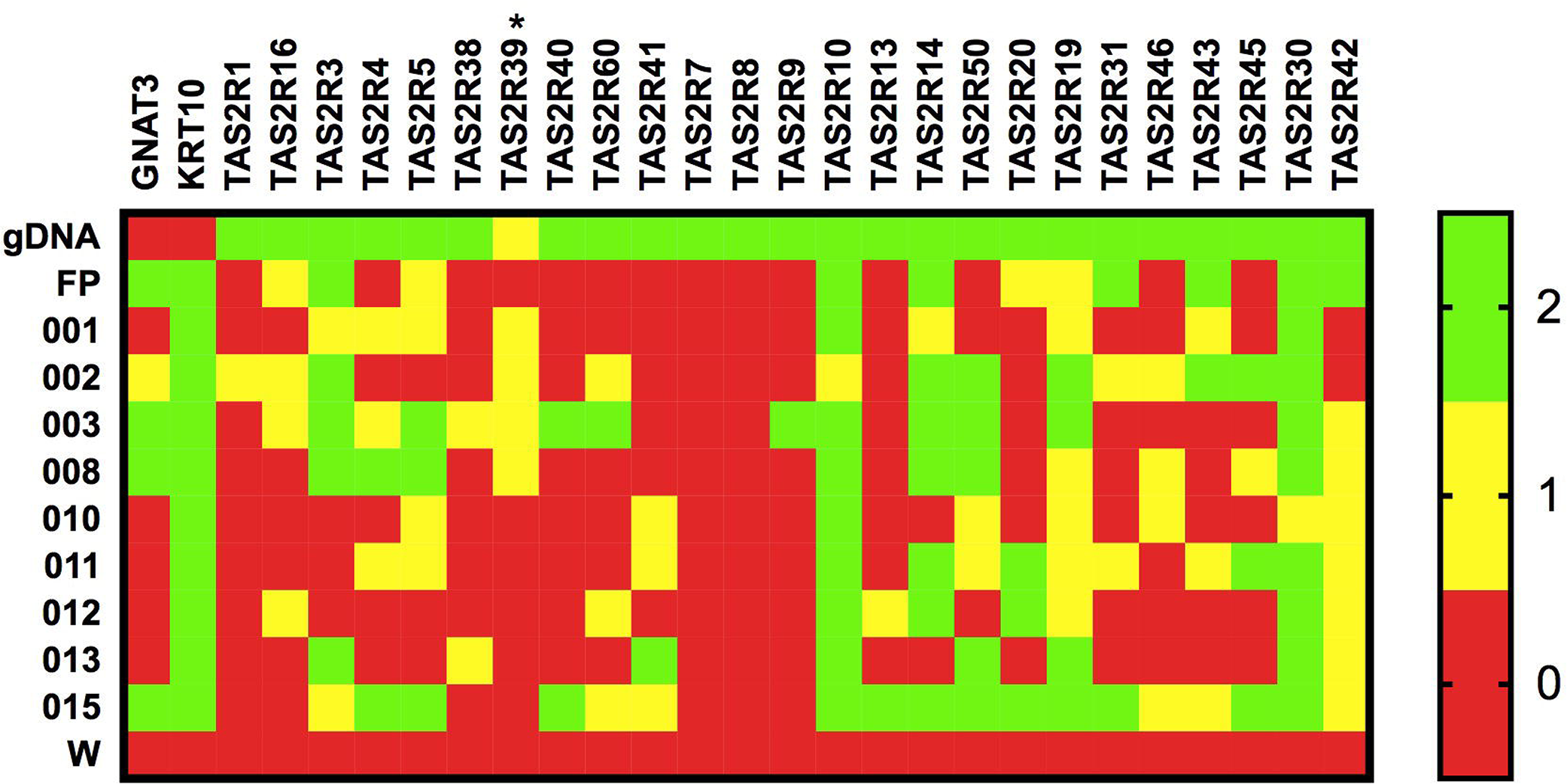
Results from two rounds of PCR. Each column is labeled by a gene, with members of the *TAS2R* family in the order of location on human chromosomes. Each row is labeled by a sample ID, where ‘gDNA’ represents genomic DNA (positive control), ‘FP’ represents taste tissue, and ‘W’ represents water (a negative control). Green box, bands in both experiments; yellow box, bands in one experiment; red box, no bands. ^*^ indicates that there was only one PCR experiment for that gene.

**Table 2:**
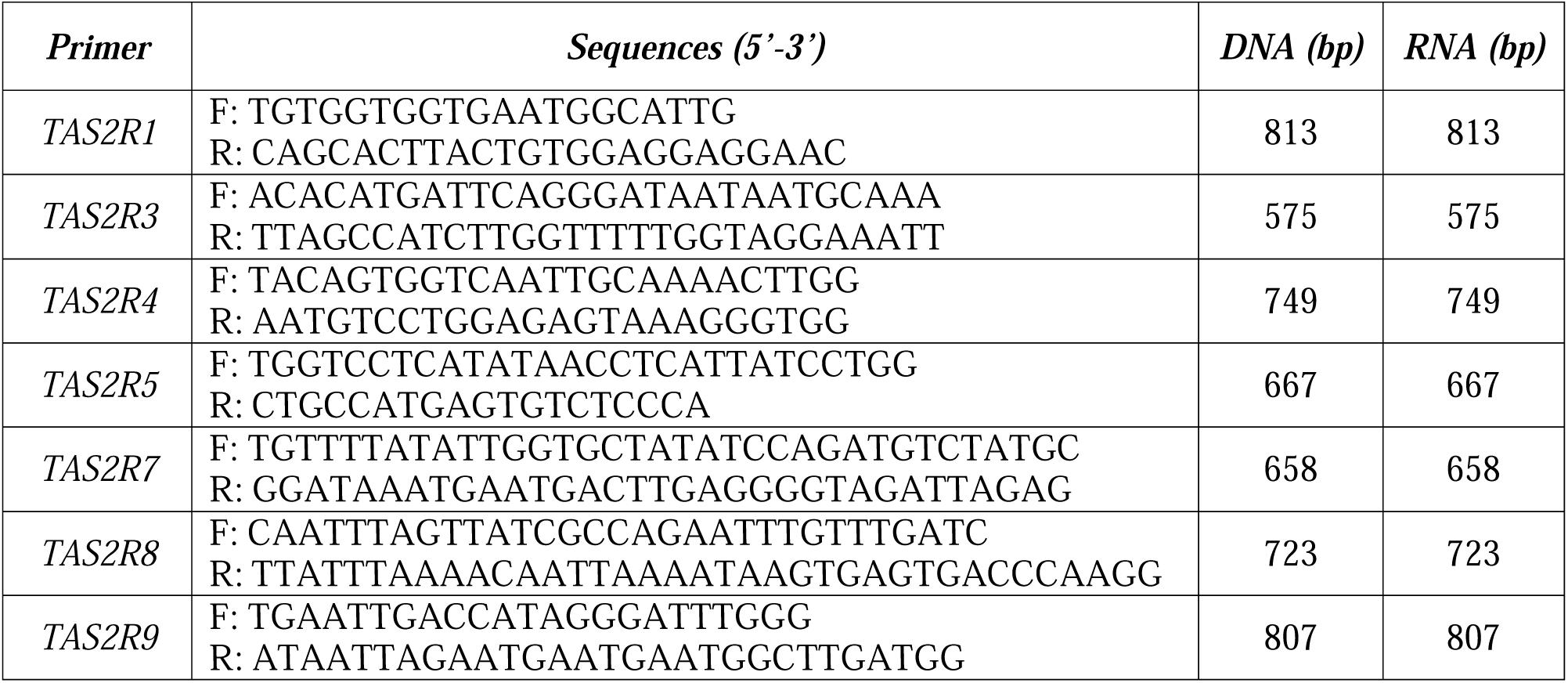
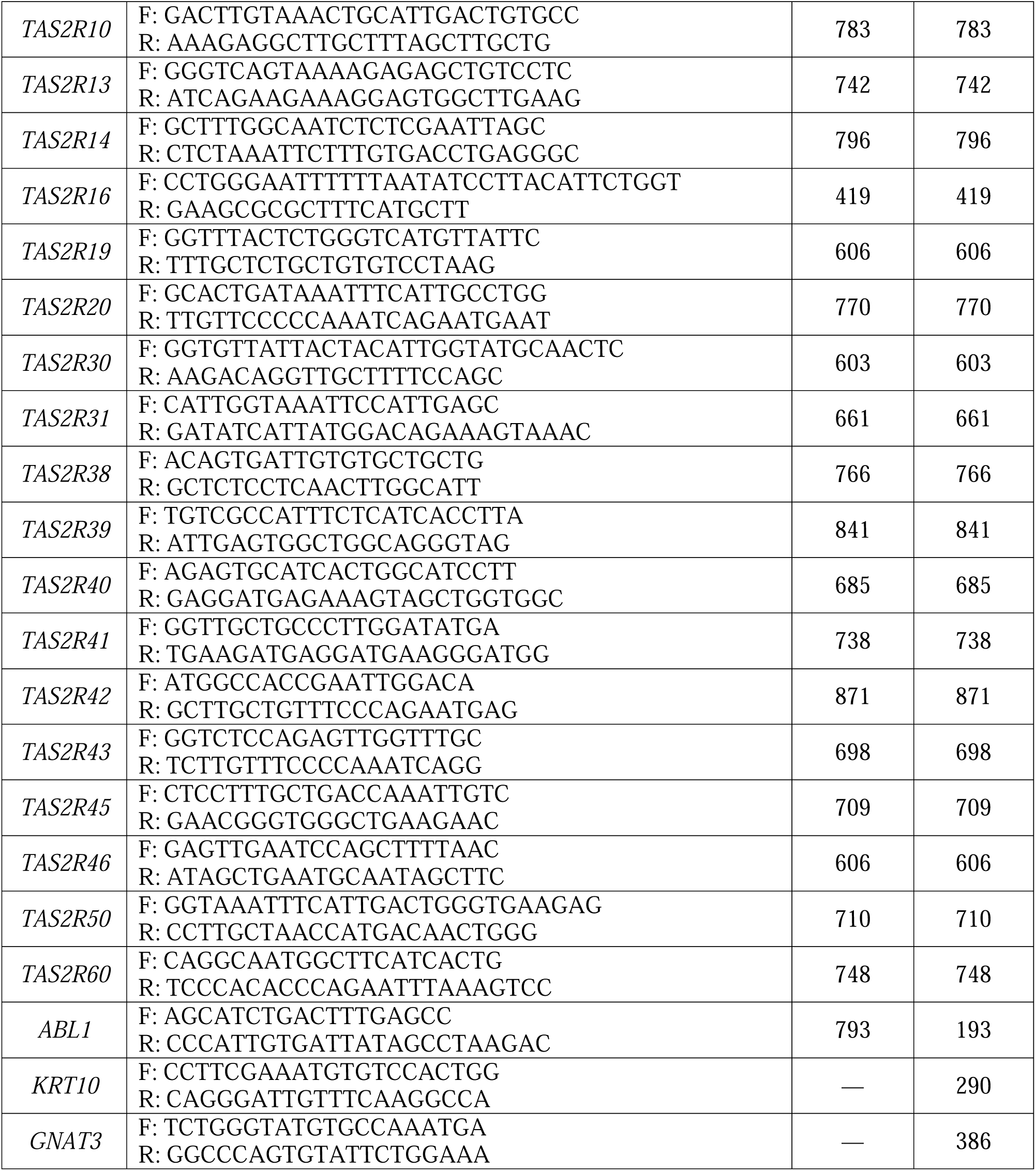
Primer sequences.

The oligonucleotide sequences and the corresponding amplicon sizes are given for genomic DNA and cDNA. F, Forward; R, reverse; bp, base pairs.

### Quantitative PCR analysis

To quantify mRNA abundance, qPCR was performed on each of the 25 *TAS2R* genes standardized to the housekeeping gene *GAPDH* (Fig 4). The results show variable expression of the *TAS2R* genes across samples, which was expected based on the results of the PCR amplification experiments. The taste tissue sample showed variable expression across receptor type. We also confirmed some expression of *GNAT3* in samples using qPCR standardized to the housekeeping gene *GAPDH* (Fig 4).

**Figure 4:**
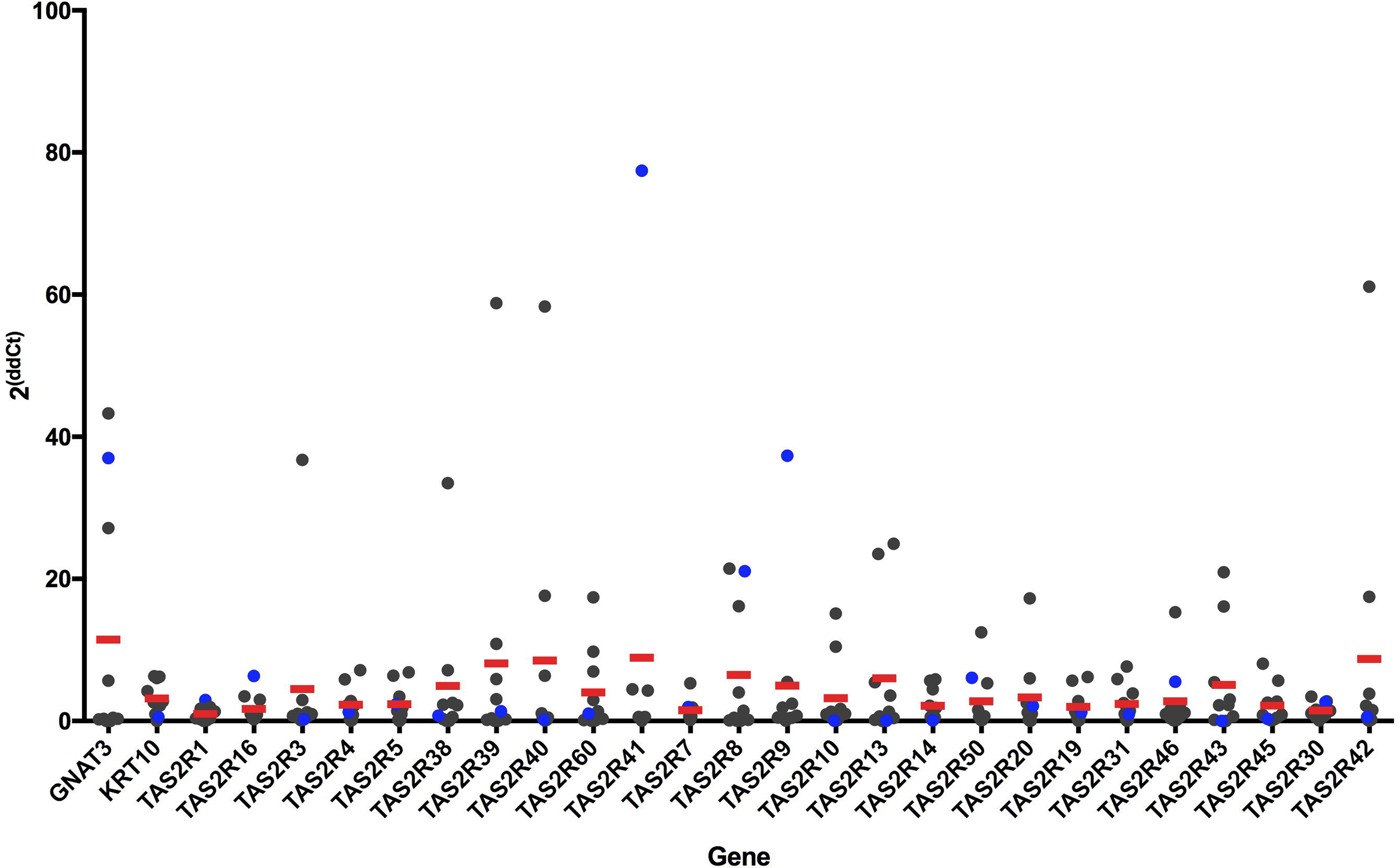
Quantification of bitter taste-related gene expression—qPCR results from cDNA of skin samples after amplification for genes of interest. cDNA was amplified with primers for *GNAT3*, *KRT10*, and the 25 *TAS2R* genes. Data were standardized to the housekeeping gene *GAPDH*, and 2^ΔΔCt^ was calculated. Results were plotted with individual values in gray and mean across all subjects in red (*n* = 9). Data points for the FP sample are in blue.

Taste-related genes are minimally expressed even in taste tissue, and the results here were variable, as is the case with expression of mRNA near the level of detection [16]. Despite these limitations, these results suggested that a study of *TAS2R* mRNA expression in skin with a larger sample size was warranted. To do so, we turned to a large and publicly available RNA-seq data set.

### GTEx data analysis

After appropriate approvals, we obtained RNA-seq expression data from the Genotype-Tissue Expression project (GTEx; #12732: Bitter receptor gene expression: patterns across tissues). The data were measured at the gene level in RPKM units (reads per kilobase of transcript per million mapped reads) and we extracted the expression data for 25 bitter receptor genes. The data analyzed consisted of 914 skin samples that varied in presumed sun exposure (sun-exposed from lower leg or not-sun-exposed from suprapubic region), sex, and age (Table 1). This data set was used because RNA-seq provides more accurate detection of low-abundance transcripts and because it provided a large sample size. There was heterogeneity of variance between the *TAS2R* genes, but the most highly expressed bitter receptor genes were *TAS2R5*, *14*, *20*, and *4* (Fig 5). For statistical analysis of sun-exposure, we considered only subjects who had donated both sun-exposed and not sun-exposed tissue (n=299) and performed Kruskal-Wallis tests to detect differences in the distribution of gene expression levels based on sun-exposure. Results based on tissue type indicated significantly lower expression levels in sun-exposed skin for *TAS2R4* (Median diff. =0.084, p < 0.05), *TAS2R30* (Median diff.=0.009, p<0.01), and *TAS2R42* (Median diff=0, p < 0.05), but significantly higher expression levels in sun-exposed skin for *TAS2R60* (Median diff=0.046, p<0.0001) (Fig 6 and S1 Table). We also observed a small sex difference in mRNA expression. In skin from the suprapubic area, females” expression was significantly higher for *TAS2R3* (Median diff. = 0.034, p < 0.01), *TAS2R4* (Median diff. = 0.126, p < 0.01), and *TAS2R8* (Median diff = 0, p < 0.05) (Fig 7, S2 Table). In skin from the lower leg, females” expression was significantly lower for *TAS2R3* (Median diff. = 0.023, p < 0.05), *TAS2R9* (Median diff = 0, p < 0.01), and *TAS2R14* (Median diff = 0.080, p < 0.01) (Fig 7, S3 Table). Finally, there was a positive correlation between increasing age and expression of *TAS2R5* gene but only in not-sun-exposed skin (*p* = 0.001) (Fig 8).

**Figure 5:**
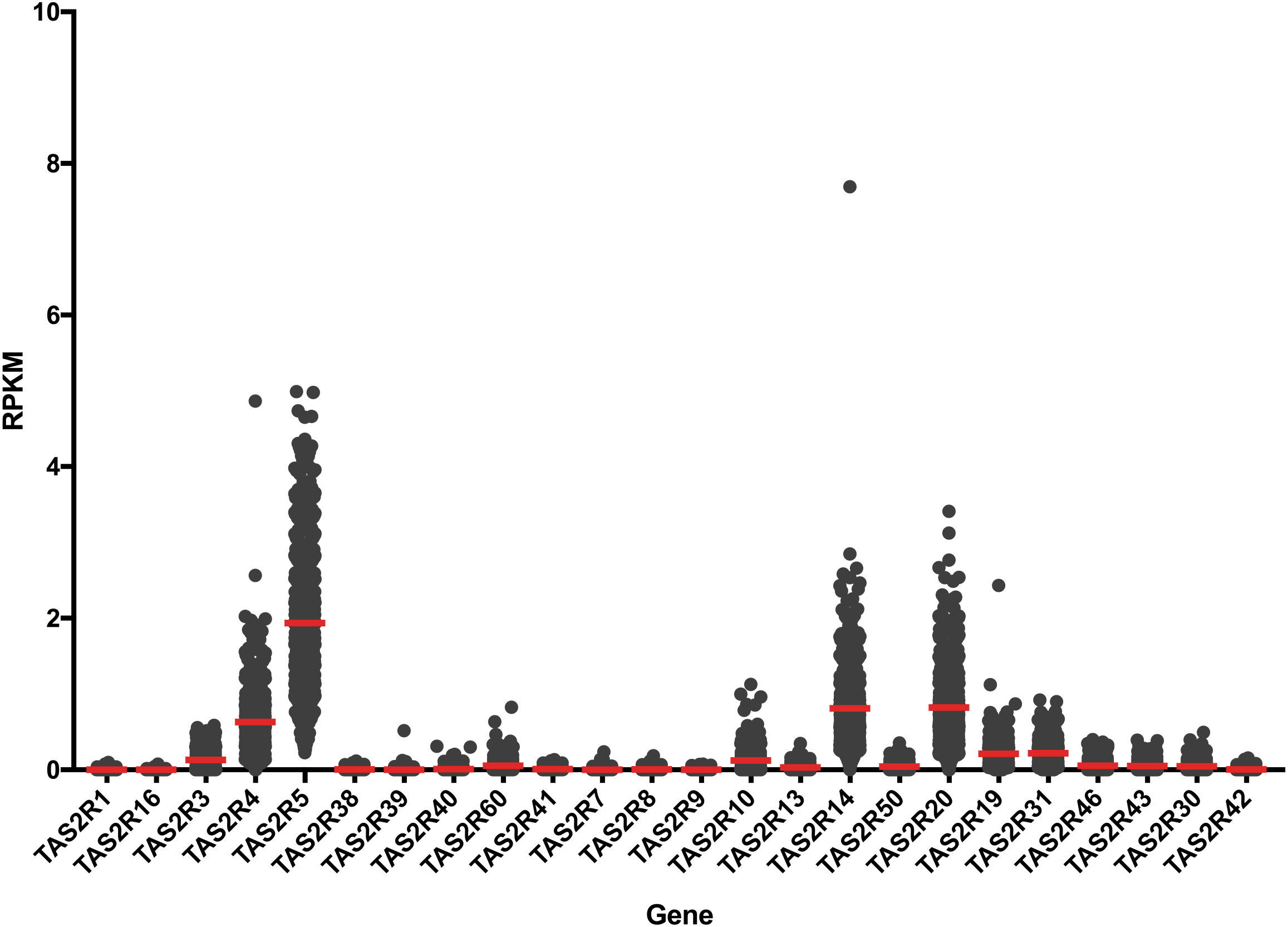
Expression levels of *TAS2R* genes from RNA-seq obtained from the GTEx database. Data are plotted with individual RPKM values in gray points and mean across all samples in red lines (*N* = 914).

**Figure 6:**
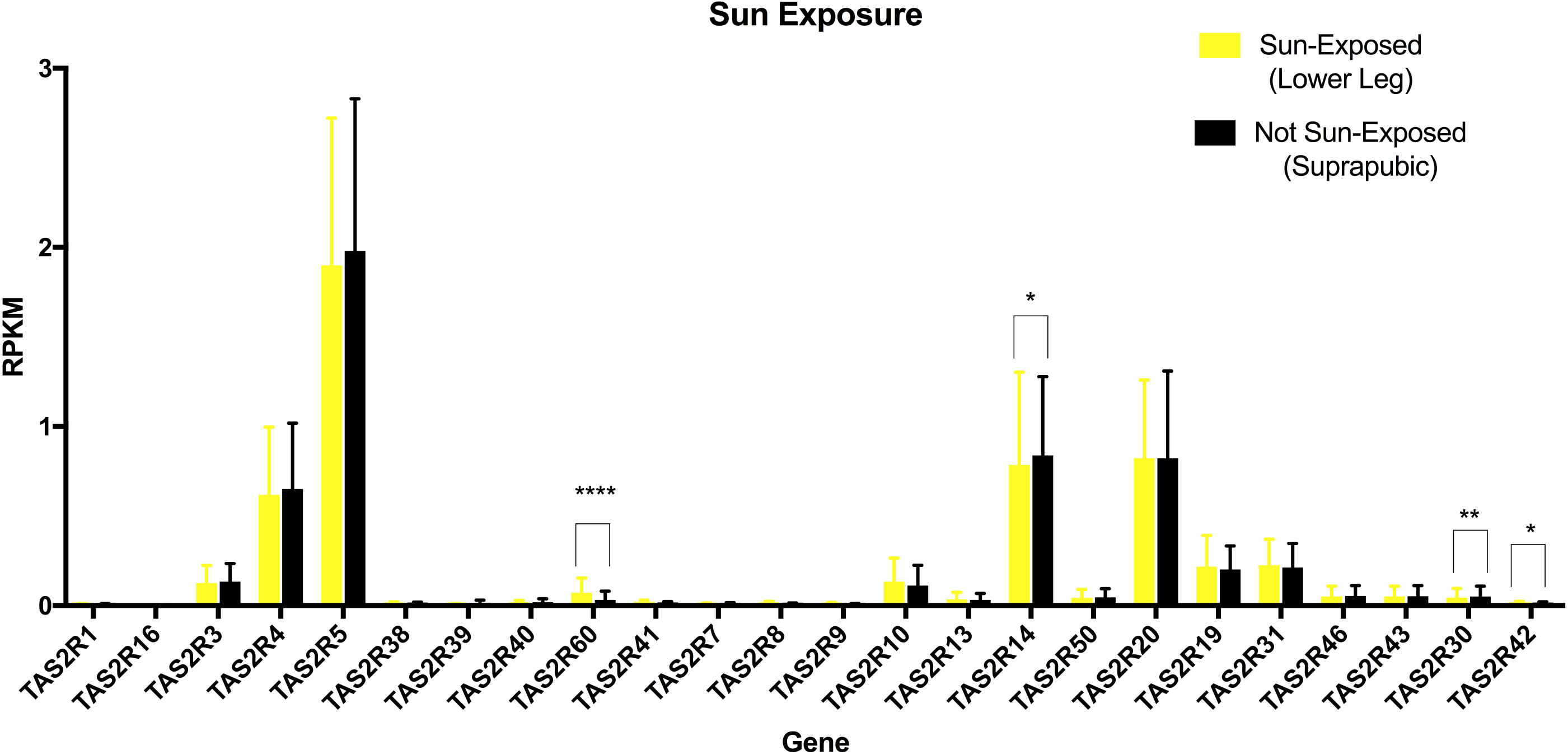
Effect of sun exposure on *TAS2R* expression from the GTEx data. Expression levels of bitter receptor genes from RNA-seq obtained from the GTEx database are separated based on sun exposure. Data are plotted as mean and SD across subjects that donated both skin sample types (N = 299 for each sample type). ^*^p<0.05, ^**^p<0.01, ^***^*p* < 0.001; ^****^*p* < 0.0001.

**Figure 7:**
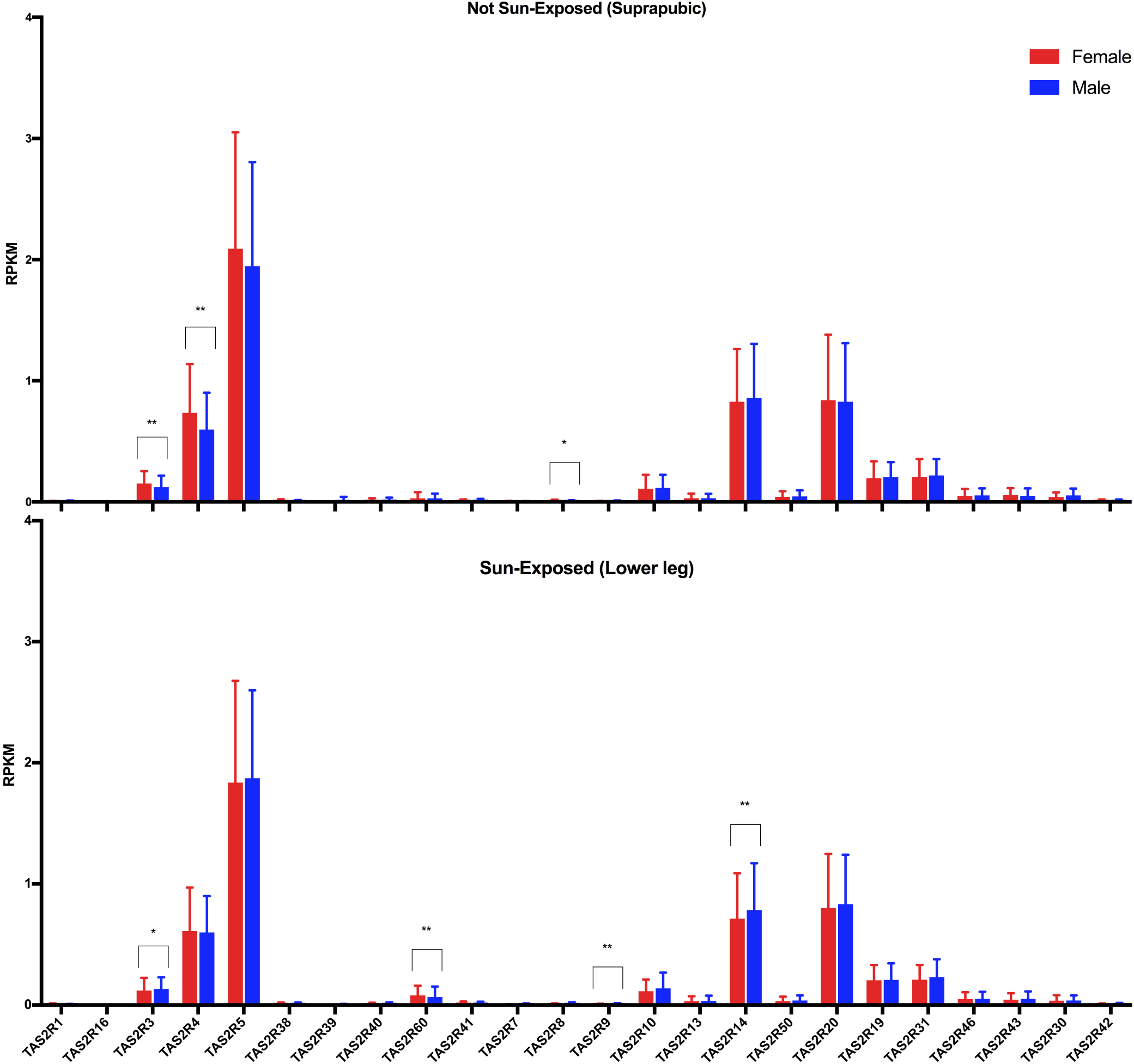
Effect of sex on *TAS2R* expression from the GTEx data. Expression levels of bitter receptor genes from RNA-seq obtained from the GTEx database are separated based on sex and presumed sun exposure. Data are plotted as mean and SD across males (N = 603) and females (N = 311) that donated both skin sample types. ^*^p<0.05, ^**^p<0.01, ^***^*p* < 0.001; ^****^*p* < 0.0001.

**Figure 8:**
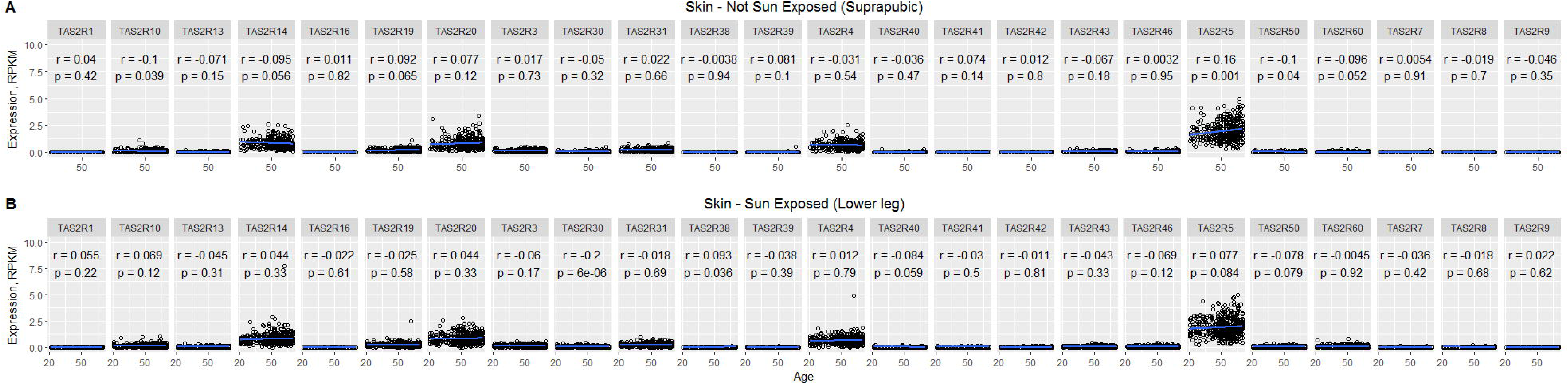
Correlation plots of *TAS2R* expression against age from the GTEx data. Individual RPKM data are plotted separated by sun exposure and in order of increasing age of the subject for each receptor. R values and p values are given on the corresponding plot.

## Discussion

Previous studies have shown bitter taste receptor expression in many tissues, including the airway, gastrointestinal tract, and testes [2]. Here, we provide a comprehensive analysis of bitter taste receptor expression in skin using two types of skin samples and three methods of analysis. This pattern of results suggests an association between *TAS2R* expression and chromosomal location. For instance, there is no expression of the *TAS2R* gene on chromosome 5 and little to no expression of the first few *TAS2Rs* on chromosome 12. We found that some bitter receptors are not expressed at all, some are variably expressed among people, and some are expressed in almost all skin samples we tested. Variability in more highly expressed receptors is related to skin location (presumed-sun-exposed vs. non-exposed), sex, and age. Expression of taste-related gene *GNAT3* suggests that these receptors are functional in the skin and that the pathway may be G protein–dependent.

The role of bitter receptors in the skin may become apparent after exploring the most highly expressed receptors and their known agonists. Some T2R proteins are promiscuous and bind to a wide variety of substances, whereas others have more specificity and bind to one or a few known substances. The protein products of *TAS2R5* and *TAS2R20*, two of the most highly expressed genes in the GTEx data set, are narrowly tuned and recognize one to three of 104 known bitter compounds [10]. T2R4, the product of *TAS2R4*, another highly expressed gene in this study, is intermediate and binds to 6–16 known bitter compounds. Finally, the *TAS2R14* product, T2R14 is broadly tuned and binds to 33 known bitter substances, including synthetic medicinal compounds [17, 18]. Interestingly, *TAS2R38*, the gene for the bitter receptor that enhances innate immunity of the upper respiratory system by recognizing bacteria [8], is rarely or never expressed in skin. We do not know whether the agonists for bitter receptors in skin are endogenous compounds, a pathogen product, or some other exogenous ligand. Further experiments should investigate the cellular response in skin when exposed to compounds similar to known agonists of these bitter receptors to learn more about their potential functions.

Determining the cellular expression of T2R proteins in skin is an important next step. Bitter receptors are typically expressed in cells known to have chemosensory functions and these cell types are typically sparsely distributed (nose, gut, and tongue). Although we do not know which cell type in human skin expresses *TAS2R* mRNA, previous studies suggest that they may be in the epidermis, and potentially expressed by keratinocytes [10, 11]. There may also be previously uncharacterized cell types in human skin similar to solitary chemosensory cells that express bitter receptors [19], where we speculate that they may function in innate immunity, wound healing, and/or differentiation. Future studies should attempt to determine the localization of T2Rs in skin potentially through immunocytochemistry, which would require validating human T2R antibodies, or *in situ* mRNA hybridization.

## Materials and Methods

### Sample collection and DNA/RNA extraction

Staff at the University of Pennsylvania Department of Dermatology collected healthy skin from 15 Mohs surgery patients for this study (n = 4 female/11 male; mean age, 62 ± 11.24 years). The Mohs procedure is used to remove cancerous skin and requires removal of additional healthy skin to facilitate proper closure of the wound [13]. We received this additional healthy skin on the day of its removal. The information obtained about each subject was provided by the department and is summarized in Table 1. Removal location was provided and based on that information as well as the proximity to cancerous skin we presumed that all samples should be considered sun-exposed. We also obtained one FP biopsy from the tongue of a separate donor as a positive control for *TAS2R* expression. FP were removed from the surface of the tongue using curved spring micro-scissors [20]. The papillae and skin tissue (0.5 mg) were mechanically homogenized and DNA and RNA was extracted using the Zymo Duet DNA/RNA MiniPrep Plus kit, following the protocol for solid tissue. DNA and RNA was quantified with the Thermo Fisher Scientific NanoDrop 1000 Spectrophotometer and measured RNA degradation through RNA integrity number equivalents generated by the Agilent TapeStation and High Sensitivity ScreenTape Assay. The RNA underwent an extra DNAse treatment using the Thermo Fisher TURBO DNA-free Kit; RNA (100ng) in water (5 μL) was then reverse transcribed into cDNA using the NuGEN Ovation RNA Amplification System V2 protocol, purified with the QIAquick PCR Purification Kit, and again quantified. The Institutional Review Board at the University of Pennsylvania approved the collection of skin biopsies for this use.

### Primers and PCR amplification

Primer sets for *KRT10* and *GNAT3* were designed using the NCBI Primer-BLAST tool. The *ABL1* primers are designed to span introns, leading to expected bands at 793 base pairs for genomic DNA and 193 base pairs for cDNA[12]. Primer sets for all 25 *TAS2R* genes have been previously published [11]. PCR reactions using primers listed in Table 2 (Invitrogen, Carlsbad, CA, USA) were performed according to the Invitrogen^™^ Platinum^™^ Taq Green Hot Start DNA Polymerase protocol with a 1 μL template. The total amount of genomic DNA from each sample was 10 ng, and the total amount of cDNA from each sample was 50 ng. A StepOne Thermocycler was used according to the following profile: one cycle of 4 min at 94 °C; 40 cycles of 1 min at 94 °C, 1 min at 55 °C, 2 min at 72 °C; one cycle of a final hold at 4 °C. Fragments were detected by staining with SYBR Green Safe. The PCR products were electrophoresed on a 1.0% gel in TAE buffer.

### Real-time qPCR

Real Time qPCR reactions were performed in 10 μL of water in a 384-well plate according to the TaqMan^®^ Fast Advanced Master Mix protocol with 1 μL template and run in triplicate. The total amount of cDNA from each sample was 50 ng. Primers for skin-specific markers, *TAS2Rs*, and a pre-developed endogenous control, *GAPDH* were used. PCR reactions were performed with the QuantStudio 12K Flex Real-Time PCR machine and amplification was evaluated by comparative analysis based on cycle threshold [21]. Graphs were generated using GraphPad Prism 7 (La Jolla, CA, USA).

### GTEx database analysis

RNA-seq data from 914 post-mortem tissue samples were provided by the GTEx project (Table 1), with information about each sample, including the age and sex of the tissue donor, and tissue type (sun-exposed skin from lower leg or sun-unexposed skin from suprapubic region). For the 25 bitter receptor genes from 914 samples, the gene expression RPKM values were normalized for all samples of the same tissue type. Due to the heterogeneity of variance between the genes, we used the non-parametric Kruskal-Wallis test to detect differences in the distribution of expression levels based on effects of sun exposure and of sex within each tissue type (S1-3 Tables). For analysis of effects of sun exposure, only data from the 299 subjects that donated both types of samples were included. For effects of sex in skin from the lower leg, all 508 tissue samples were included, and from the suprapubic area, all 406 tissue samples were included. Data for sun exposure effects and sex were analyzed in R version 3.4.2, and graphs were generated in GraphPad Prism 7. Effects of age were analyzed via correlation and plotted in R (version 3.4.2) and R-studio (version 1.0.136). We deposited a data analysis script based in R on Github (https://github.com/DanielleReed/TAS2R38).

## Acknowledgments

Dr. Aimee Payne and the staff at the University of Pennsylvania Department of Dermatology are acknowledged for providing samples and technical assistance. We are thankful to Dr. Casey Trimmer for sharing her expertise and providing advice for the study. Dr. Mary Matsui, and Dr. Ed Pelle from the Estée Lauder Companies are also acknowledged for their contribution to our preliminary work and for their insight on the study. We are grateful to Nora Ruth from The Estée Lauder Companies for revising the manuscript.

## Supporting Information

**S1 Figure: Gene expression of *ABL1***. PCR was performed with genomic DNA from skin (gDNA), a mixture of genomic DNA and cDNA from skin (Mix), cDNA from fungiform papillae (FP), and cDNA from 14 skin samples (001-015). Water was used as a no-template control. The larger band at 793 base pairs (bp) includes introns, and the smaller band at 293 bp does not contain introns. Genomic DNA was used as a positive control for the larger band size. A mix was used as a positive control for both bands. The smear at FP is likely caused by nonspecific binding.

**S2 Figure: Gene expression of *TAS2R1***. PCR was performed with genomic DNA from skin (gDNA), cDNA from fungiform papillae (FP), and cDNA from nine skin samples. Water was used as a no-template control. The expected band size is 813 bp. The experiment was replicated (bottom panel) because taste receptors are not abundant and can have variable results.

**S3 Figure: Gene expression of *TAS2R3***. PCR was performed with genomic DNA from skin (gDNA), cDNA from fungiform papillae (FP), and cDNA from nine skin samples. Water was used as a no-template control. The expected band size is 575 bp. The experiment was replicated (bottom panel) because taste receptors are not abundant and can have variable results.

**S4 Figure: Gene expression of *TAS2R4***. PCR was performed with genomic DNA from skin (gDNA), cDNA from fungiform papillae (FP), and cDNA from nine skin samples. Water was used as a no-template control. The expected band size is 749 bp. The experiment was replicated (bottom panel) because taste receptors are not abundant and can have variable results.

**S5 Figure: Gene expression of *TAS2R5***. PCR was performed with genomic DNA from skin (gDNA), cDNA from fungiform papillae (FP), and cDNA from nine skin samples. Water was used as a no-template control. The expected band size is 667 bp. The experiment was replicated (bottom panel) because taste receptors are not abundant and can have variable results.

**S6 Figure: Gene expression of *TAS2R7***. PCR was performed with genomic DNA from skin (gDNA), cDNA from fungiform papillae (FP), and cDNA from nine skin samples. Water was used as a no-template control. The expected band size is 658 bp. The experiment was replicated (bottom panel) because taste receptors are not abundant and can have variable results.

**S7 Figure: Gene expression of *TAS2R8***. PCR was performed with genomic DNA from skin (gDNA), cDNA from fungiform papillae (FP), and cDNA from nine skin samples. Water was used as a no-template control. The expected band size is 723 bp. The experiment was replicated (bottom panel) because taste receptors are not abundant and can have variable results.

**S8 Figure: Gene expression of *TAS2R9***. PCR was performed with genomic DNA from skin (gDNA), cDNA from fungiform papillae (FP), and cDNA from nine skin samples. Water was used as a no-template control. The expected band size is 807 bp. The experiment was replicated (bottom panel) because taste receptors are not abundant and can have variable results.

**S9 Figure: Gene expression of *TAS2R10***. PCR was performed with genomic DNA from skin (gDNA), cDNA from fungiform papillae (FP), and cDNA from nine skin samples. Water was used as a no-template control. The expected band size is 783 bp. The experiment was replicated (bottom panel) because taste receptors are not abundant and can have variable results.

**S10 Figure: Gene expression of *TAS2R13***. PCR was performed with genomic DNA from skin (gDNA), cDNA from fungiform papillae (FP), and cDNA from nine skin samples. Water was used as a no-template control. The expected band size is 742 bp. The experiment was replicated (bottom panel) because taste receptors are not abundant and can have variable results.

**S11 Figure: Gene expression of *TAS2R14***. PCR was performed with genomic DNA from skin (gDNA), cDNA from fungiform papillae (FP), and cDNA from nine skin samples. Water was used as a no-template control. The expected band size is 796 bp. The experiment was replicated (bottom panel) because taste receptors are not abundant and can have variable results.

**S12 Figure: Gene expression of *TAS2R16***. PCR was performed with genomic DNA from skin (gDNA), cDNA from fungiform papillae (FP), and cDNA from nine skin samples. Water was used as a no-template control. The expected band size is 419 bp. The experiment was replicated (bottom panel) because taste receptors are not abundant and can have variable results.

**S13 Figure: Gene expression of *TAS2R19***. PCR was performed with genomic DNA from skin (gDNA), cDNA from fungiform papillae (FP), and cDNA from nine skin samples. Water was used as a no-template control. The expected band size is 606 bp. The experiment was replicated (bottom panel) because taste receptors are not abundant and can have variable results.

**S14 Figure: Gene expression of *TAS2R20***. PCR was performed with genomic DNA from skin (gDNA), cDNA from fungiform papillae (FP), and cDNA from nine skin samples. Water was used as a no-template control. The expected band size is 770 bp. The experiment was replicated (bottom panel) because taste receptors are not abundant and can have variable results.

**S15 Figure: Gene expression of *TAS2R30***. PCR was performed with genomic DNA from skin (gDNA), cDNA from fungiform papillae (FP), and cDNA from nine skin samples. Water was used as a no-template control. The expected band size is 603 bp. The experiment was replicated (bottom panel) because taste receptors are not abundant and can have variable results.

**S16 Figure: Gene expression of *TAS2R31***. PCR was performed with genomic DNA from skin (gDNA), cDNA from fungiform papillae (FP), and cDNA from nine skin samples. Water was used as a no-template control. The expected band size is 661 bp. The experiment was replicated (bottom panel) because taste receptors are not abundant and can have variable results.

**S17 Figure: Gene expression of *TAS2R38***. PCR was performed with genomic DNA from skin (gDNA), cDNA from fungiform papillae (FP), and cDNA from nine skin samples. Water was used as a no-template control. The expected band size is 766 bp. Multiple bands are likely because of non-specific binding. The experiment was replicated (bottom panel) because taste receptors are not abundant and can have variable results.

**S18 Figure: Gene expression of *TAS2R39***. PCR was performed with genomic DNA from skin (gDNA), cDNA from fungiform papillae (FP), and cDNA from nine skin samples. Water was used as a no-template control. The expected band size is 841 bp. The experiment was replicated, but results were omitted because of non-specific binding.

**S19 Figure: Gene expression of *TAS2R40***. PCR was performed with genomic DNA from skin (gDNA), cDNA from fungiform papillae (FP), and cDNA from nine skin samples. Water was used as a no-template control. The expected band size is 685 bp. The experiment was replicated (bottom panel) because taste receptors are not abundant and can have variable results.

**S20 Figure: Gene expression of *TAS2R41***. PCR was performed with genomic DNA from skin (gDNA), cDNA from fungiform papillae (FP), and cDNA from nine skin samples. Water was used as a no-template control. The expected band size is 738 bp. Multiple bands are likely because of non-specific binding. The experiment was replicated (bottom panel) because taste receptors are not abundant and can have variable results.

**S21 Figure: Gene expression of *TAS2R42***. PCR was performed with genomic DNA from skin (gDNA), cDNA from fungiform papillae (FP), and cDNA from nine skin samples. Water was used as a no-template control. The expected band size is 871 bp. The experiment was replicated (bottom panel) because taste receptors are not abundant and can have variable results.

**S22 Figure: Gene expression of *TAS2R43***. PCR was performed with genomic DNA from skin (gDNA), cDNA from fungiform papillae (FP), and cDNA from nine skin samples. Water was used as a no-template control. The expected band size is 698 bp. The experiment was replicated (bottom panel) because taste receptors are not abundant and can have variable results.

**S23 Figure: Gene expression of *TAS2R45***. PCR was performed with genomic DNA from skin (gDNA), cDNA from fungiform papillae (FP), and cDNA from nine skin samples. Water was used as a no-template control. The expected band size is 709 bp. Multiple bands are likely because of non-specific binding. The experiment was replicated (bottom panel) because taste receptors are not abundant and can have variable results.

**S24 Figure: Gene expression of *TAS2R46***. PCR was performed with genomic DNA from skin (gDNA), cDNA from fungiform papillae (FP), and cDNA from nine skin samples. Water was used as a no-template control. The expected band size is 606 bp. The experiment was replicated (bottom panel) because taste receptors are not abundant and can have variable results.

**S25 Figure: Gene expression of *TAS2R50***. PCR was performed with genomic DNA from skin (gDNA), cDNA from fungiform papillae (FP), and cDNA from nine skin samples. Water was used as a no-template control. The expected band size is 710 bp. The experiment was replicated (bottom panel) because taste receptors are not abundant and can have variable results.

**S26 Figure: Gene expression of *TAS2R60***. PCR was performed with genomic DNA from skin (gDNA), cDNA from fungiform papillae (FP), and cDNA from nine skin samples. Water was used as a no-template control. The expected band size is 748 bp. The experiment was replicated (bottom panel) because taste receptors are not abundant and can have variable results.

**S27 Figure: Gene expression of *GNAT3***. PCR was performed with genomic DNA from skin (gDNA), cDNA from fungiform papillae (FP), and cDNA from nine skin samples. Water was used as a no-template control. The primer set is intron-spanning, so there is no expected band size for genomic DNA, while there is an expected band size of 386 bp for cDNA. The experiment was replicated (bottom panel) because taste receptors are not abundant and can have variable results.

**S28 Figure: Gene expression of *KRT10***. PCR was performed with genomic DNA from skin (gDNA), cDNA from fungiform papillae (FP), and cDNA from nine skin samples. Water was used as a no-template control. The primer set is intron-spanning, so there is no expected band size for genomic DNA and an expected band size of 290 bp for cDNA. The experiment was replicated (bottom panel) because taste receptors are not abundant and can have variable results.

**S1 Table**. Kruskal-Wallis test statistics for GTEx data comparing effects of presumed sun exposure for each gene of interest (N=598).

**S2 Table**. Kruskal-Wallis test statistics for GTEx data comparing effects of sex for each gene of interest in not sun-exposed tissue (N=406).

**S3 Table**. Kruskal-Wallis test statistics for GTEx data comparing effects of sex for each gene of interest in sun-exposed tissue (N=508).

